# Detecting fabrication in large-scale molecular omics data

**DOI:** 10.1101/757070

**Authors:** Michael S. Bradshaw, Samuel H. Payne

**Affiliations:** Computer Science Department, University of Colorado Boulder, Boulder CO 80309 USA; Biology Department, Brigham Young University, Provo UT 84602 USA

## Abstract

Fraud is a pervasive problem and can occur as fabrication, falsification, plagiarism or theft. The scientific community is not exempt from this universal problem and several studies have recently been caught manipulating or fabricating data. Current measures to prevent and deter scientific misconduct come in the form of the peer-review process and on-site clinical trial auditors. As recent advances in high-throughput omics technologies have moved biology into the realm of big-data, fraud detection methods must be updated for sophisticated computational fraud. In the financial sector, machine learning and digit-preference are successfully used to detect fraud. Drawing from these sources, we develop methods of fabrication detection in biomedical research and show that machine learning can be used to detect fraud in large-scale omic experiments. Using the raw data as input, the best machine learning models correctly predicted fraud with 84-95% accuracy. With digit frequency as input features, the best models detected fraud with 98%-100% accuracy. All of the data and analysis scripts used in this project are available at https://github.com/MSBradshaw/FakeData.

## Introduction

Fraud is a pervasive problem and can occur as fabrication, falsification, plagiarism or theft. Examples of fraud are found in virtually every field, such as: education, commerce and technology. With the rise of electronic crimes, specific criminal justice and regulatory bodies have been formed to detect sophisticated fraud, creating an arms-race between methods to deceive and methods to detect deception. The scientific community is not exempt from the universal problem of fraud, and several studies have recently been caught manipulating or fabricating data [1,2] or are suspected of it [3]. More than two million scientific articles are published yearly and ~2% of authors admit to data fabrication [4]. When asked if their colleagues had fabricated data, positive response rates rose to 14-19% [4,5]. Some domains or locales have somewhat higher rates of data fabrication; in a recent survey of researchers at Chinese hospitals, 7.37% of researchers admitted to fabricating data [6]. Overall, these rates of data fabrication potentially means tens to hundreds of thousands of articles are published each year with manipulated data.

Data in the biological sciences is particularly vulnerable to fraud given its size - which makes it easier to hide data manipulation - and researcher’s dependence on freely available public data. Recent advances in high-throughput omics technologies have moved biology into the realm of big-data. Many diseases are now characterized in populations, with thousands of individuals characterized for cancer [7], diabetes [8], bone strength [9], and health care services for the general populace [10]. Large-scale characterization studies are also done for cell lines and drug responses [11,12]. With the rise of importance of these large datasets, it becomes imperative that they remain free of errors both unintentional and intentional [13].

Current methods for ensuring the validity of research is largely limited to the peer-review process which as of late has proven to be insufficient at spotting blatant duplication of images [14], let alone subtleties hidden in large scale data. Data for clinical trials can be subject to reviews and central monitoring [15,16]. However, the decision regarding oversight methodology and frequency is not driven by empirical data, but rather is determined by clinics’ usual practice [17]. The emerging data deluge challenges the effectiveness of traditional auditing practices to detect fraud, and several studies have suggested addressing the issue with improved centralized and independent statistical monitoring [5,6,16,18]. However, these recommendations are given chiefly to help ensure the safety and efficacy of the study, not data integrity.

In 1937, physicist Frank Benford observed in a compilation of 20,000 numbers that the first digit did not follow a uniform distribution as one may anticipate [19]. This pattern holds true in most large collections of numbers, including scientific data. Comparing a distribution of first digits to a Benford distribution can be used to identify deviations from the expected frequency, often because of fraud. Recently Benford’s law has been used to identify fraud in financial records of international trade [20] and money laundering [21]. It has also been used on a smaller scale to reaffirm suspicions of fraud in clinical trials [3].

The distinction between fraud and honest error is important to make. Fraud is the intent to cheat [5]. This is the definition used throughout this paper. An honest error might be, forgetting to include a few samples, while intentionally excluding samples would be fraud. Copying and pasting values from one table to another incorrectly is an honest error but intentionally changing the values is fraud. In these examples the results may be the same but the intent behind them differs wildly. In efforts to maintain data integrity, identifying the intent of the misconduct may be impossible, and is also a secondary consideration after suspect data has been identified.

Data fabrication is “making up data or results and recording or reporting them” [5]. This type of data manipulation is free from the above ambiguity relating to the author’s intent. Making up data is always wrong. We explore methods of data fabrication and detection in molecular omics data using supervised machine learning and Benford-like digit-frequencies. We do not attempt to explain why someone may choose to fabricate their data - as other study have done [6,22]; our only goal is to evaluate the utility of digit-frequencies to differentiate real from fake data.The data used in this study comes from the Clinical Proteomic Tumor Analysis Consortium (CPTAC) cohort for endometrial carcinoma, which contains copy number alteration (CNA) measurements from 100 tumor samples. We created 50 additional fake samples for these datasets. Three different methods of varying sophistication are used for fabrication: random number generation, resampling with replacement and imputation. We show that machine learning and digit-preference can be used to detect fraud with near perfect accuracy.

## Methods

### Real Data

The real data used in this publication originated from the genomic analysis of uterine endometrial cancer. As part of the Clinical Proteomics Tumor Analysis Consortium (CPTAC), 100 tumor samples underwent whole genome and whole exome sequencing and subsequent copy number analysis. We used the results of the copy number analysis *as is*, which is stored in our GitHub repository at https://github.com/MSBradshaw/FakeData.

### Fake Data

Fake data used in this study was generated using three different methods. In each method, we created 50 fake samples which were combined with the 100 real samples to form a mixed dataset. The first method to generate fake data was random number generation. For every gene locus, we first find the maximum and minimum values observed in the original data. A new sample is then fabricated by randomly picking a value within this gene specific range. The second method to generate fake data was sampling with replacement. For this, we create lists of all observed values across the cohort for each gene. A fake sample is created by randomly sampling from these lists with replacement. The third method to generate fake data was imputation. The R package missForrest [23] was repurposed for data fabrication. A fake sample was generated by first creating a copy of a real sample. Then we iteratively nullified 10% of the data and imputed these NAs with missForrest until every value has been imputed. See Supplemental Figure 1.

### Machine Learning Training

With a mixed dataset containing 100 real samples and 50 fake samples, we proceeded to create and evaluate machine learning models which predict whether a sample is real or fabricated (Supplemental Figure 2). The 100 real and 50 fake samples were both randomly split in half, one portion added to a training set and the other held out for testing. Using Python’s SciKitLearn library, we evaluated multiple machine learning models, gradient boosting (GBD), Naïve Bayes, Random Forest, K-Nearest Neighbor (KNN), Multi-layer Perceptron (MLP) and Support Vector Machine (SVM). Training validation was done using 10-fold cross validation. We note explicitly that the training routine was never able to use testing data. After all training was complete, the held-out test set was then fed to each model for prediction and scoring. We used simple accuracy as a metric. For each sample in the test set, ML models would predict whether it was real or fabricated. Model accuracy was calculated as the number of correct predictions divided by the number of total predictions. The entire process of fake data generation and ML training/testing was repeated 50 times. Different random seeds were used when generating each set of fake data. Thus fake samples in all 50 iterations are distinct from each other. All of the data and analysis scripts used in this project are available at https://github.com/MSBradshaw/FakeData.

### Benford-Like Digit Preferences

Benford’s Law or the first digit law has been instrumental at catching fraud in various financial situations [20,21] and in small scale clinical trials [3]. The method presented here is designed with the potential to generalize and be applied to multiple sets of data of varying types and configurations (e.i. different measured variables (features) and different quantities of variables). Machine learning typically cannot handle data where the features are not consistent in number and type. Converting all measured variables to digit frequencies circumvents this problem. Digit frequencies are calculated as the number of occurrences of a single digit (0-9) divided by the total number of features. In the method described in this paper, a sample’s features are all converted to digit frequencies of the first and second digit after the decimal. Thus for each sample the features are converted from ~17,000 copy number alterations to 20 digit preferences. Using this approach, whether a sample has 100 or 17,000 features it can still be trained on and classified by the same model.

## Results

Our goal is to explore the ability of machine learning methods to identify fabricated data hidden within large datasets. Our results do not focus on the motivations to fabricate data, nor do they explore in depth the infinite methodological ways to do so. Our study focuses on whether machine learning can be trained to correctly identify fabricated data. Our general workflow is to take real data and mix in fabricated data. When training, the machine learning model is given access to the label (i.e. real or fabricated); the model is tested or evaluated by predicting the label of data which was held back from training (see Methods).

### Fake Data

The real data used in this study comes from the Clinical Proteomic Tumor Analysis Consortium (CPTAC) cohort for endometrial carcinoma, specifically the copy number alteration (CNA) data. The form of this real data is a large table of floating point values. Rows represent individual tumor samples and columns represent genes; values in the cells are thus the copy number quantification for a single gene in an individual tumor sample. This real data was paired with fabricated data and used as an input to machine learning classification models (see Methods). Three different methods of data fabrication were used in this study: random number generation, resampling with replacement, and imputation (Supplemental Figure 1). The three methods represent three realistic ways that an unscrupulous scientist might create novel data. Each method has benefits and disadvantages, with imputation being both the most sophisticated and also the most computationally intense and complex. As seen in Figure 1, the random data clusters far from the real data. Both the resampled and imputed data cluster tightly with the real data in a PCA plot, with the imputed data also generating a few reasonable outlier samples.

**Figure 1 -.**
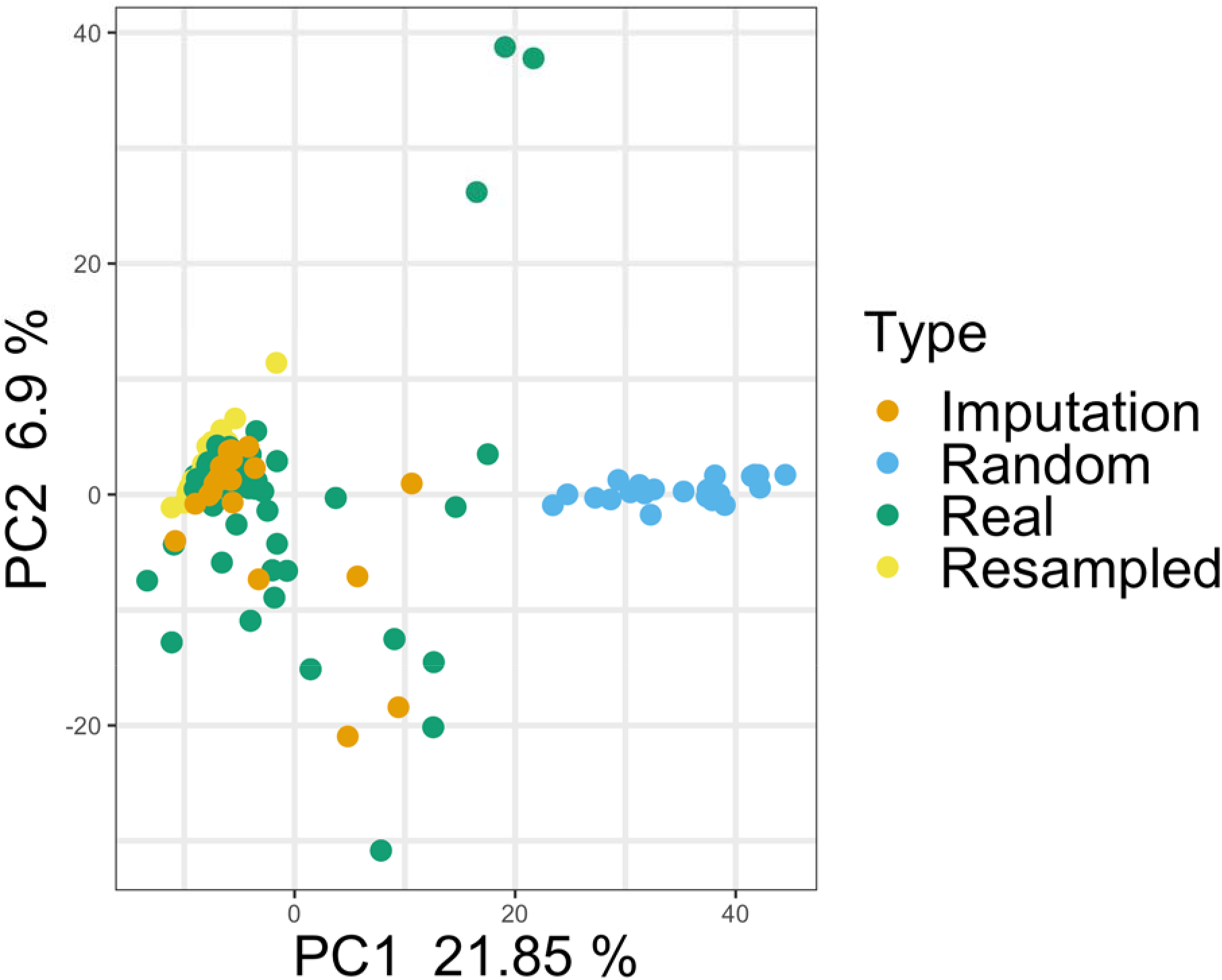
Principal Component Analysis of real and fake samples. Copy number data for the real and fabricated samples are shown. The fabricated data created via random number generation is clearly distinct from all other data. Fabricated data created via resampling or imputation appears to cluster very closely with the real data.

To look further into the fabricated data, we examined whether fake data preserved correlative relationships present in the original data (Supplemental Figure 3). This is exemplified by two pairs of genes. *PLEKHN1* and *HES4* are adjacent genes found on chromosome 1p36 separated by ~30,000 bp. Because they are so closely located on the chromosome, it is expected that most copy number events like large scale duplications and deletions would include both genes.

As expected, their CNA data has a Spearman correlation coefficient of 1.0 in the original data, a perfect correlation. The second pair of genes, *DFFB* and *OR4F5*, are also on chromosome 1, but are separated by 3.8 Mbp. As somewhat closely located genes, we would expect a modest correlation between CNA measurements, but not as highly correlated as the adjacent gene pair. Consistent with this expectation, their CNA data has a Spearman correlation coefficient of 0.27. Depending on the method of fabrication, fake data for these two gene pairs may preserve these correlative relationships. When we look at the random and resampled data for these two genes, all correlation is lost (Supplemental Figure 3 C, D, E and F). Imputation, however, produces data that closely matches the original correlations, *PLEKHN1* and *HES4* R^2^ = 0.97; *DFFB* and *OR4F5* R^2^ = 0.32 (Supplemental Figure 3 G and H).

### Machine learning with quantitative data

We tested six different methods for machine learning to create a model capable of detecting fabricated data: Gradient Boosting (GBC), Naïve Bayes, Random Forest, K-Nearest Neighbor (KNN), Multi-layer Perceptron (MLP) and Support Vector Machine (SVM). Models were given as features the quantitative data table containing copy number data on 75 labeled samples, 50 real and 25 fake. In the copy number data, each sample had measurements for ~17,000 genes, meaning that each sample had ~17,000 features. After training, the model was asked to classify held-out testing data containing 75 samples, 50 real and 25 fake. The classification task considers each sample separately, meaning that the declaration of real or fake is made only from data of a single sample. We evaluated the model on simple accuracy, whether the predicted label was correct or incorrect. To ensure that our results represent robust performance, model training and evaluation was performed 50 times; each time a completely new set of 25 fabricated samples were made (see Methods). Reported results represent the average accuracy of these 50 trials. We note that two methods, SVM and MLP, performed poorly compared to other classification methods. Testing data consisted of 2/3 real data and 1/3 fake data; therefore, baseline accuracy (the accuracy achieved if the model predicting all test samples as the majority class) is 66%. Both SVM and MLP had an average accuracy at or below this baseline for classification of the simplest fabrication method (random), and were excluded from further analysis.

The remaining four models performed relatively well on the classification task for data fabricated with the random approach. The average accuracy of 50 trials was: Random Forest 94%, GBC 92%, Naïve Bayes 88%, and KNN 72% (Figure 2A). Mean classification accuracies were lower for data created with the resampling method, with most models losing ~10% accuracy (Random Forest 84%, GBC 83%, Naïve Bayes 73%, and KNN 70%). We also note that the variability in model performance was much higher for classification of the resampled data (Figure 2B). As the resampling method uses data values from the real data, it is possible that fake samples sometimes more closely resemble real samples. Imputation classification results fluctuated (Random Forest 90%, GBC 89%, Naïve Bayes 66%, and KNN 56%). While Random Forest and GBC both increased in accuracy compared to the resampled data, Naïve Bayes and KNN both now perform at or below the baseline accuracy (Figure 2C).

**Figure 2 -.**
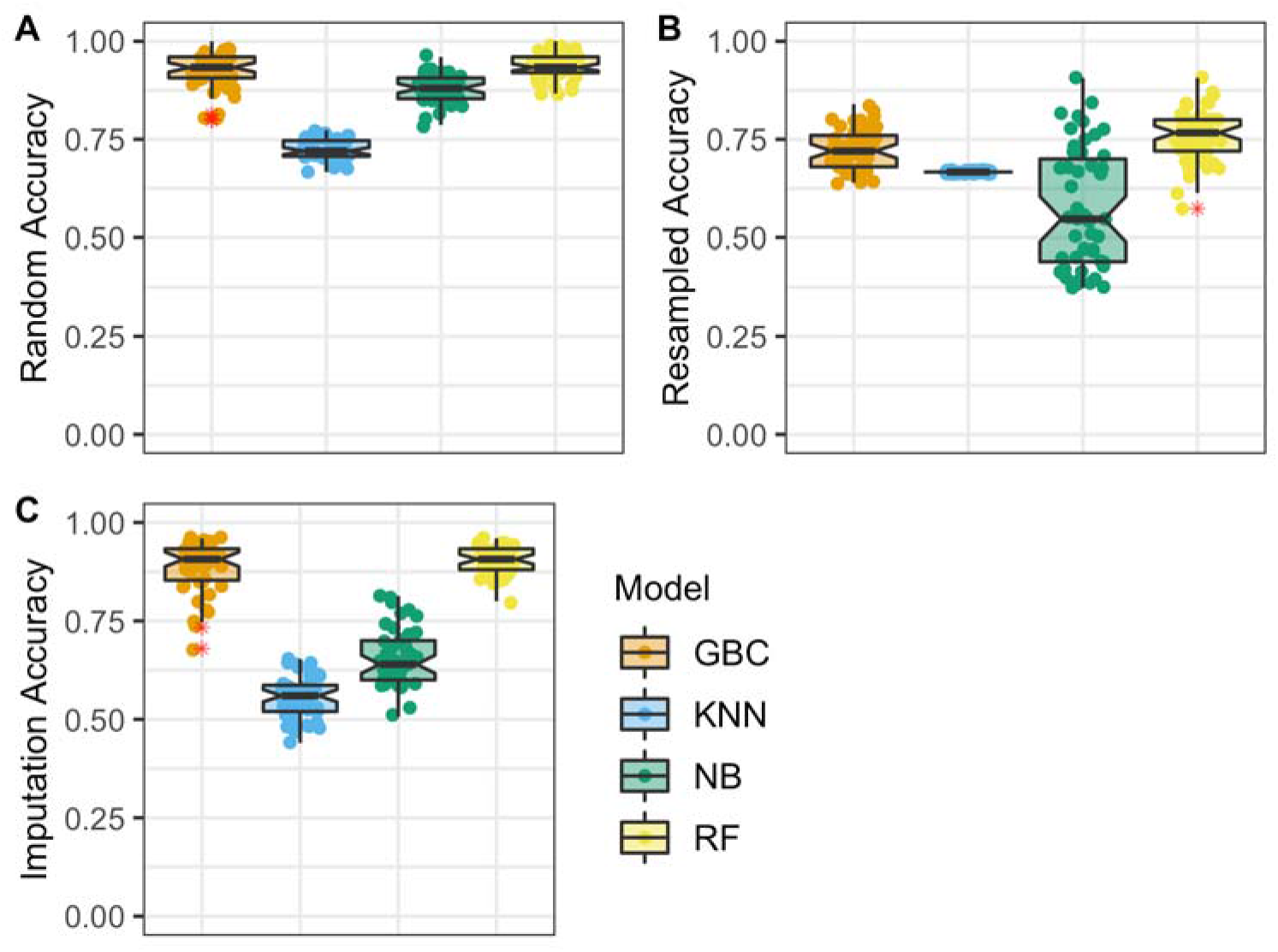
Classification accuracy using copy number data. Fabricated data was mixed with real data and given to four machine learning models for classification. Data shown represents 50 trials for 50 different fabricated dataset mixes. Features in this dataset are the copy number values for each sample. **A.** Results for data fabricated with the random method, mean classification accuracy: Random Forest 94% (+/- 3.1%), GBC 92% (+/- 4.5%), Naïve Bayes 88% (+/- 3.5%), and KNN 72% (+/- 2.6%). **B.** Results for data fabricated with the resampling method, mean classification accuracy: Random Forest 84% (+/- 6.5%), GBC 83% (+/- 5.2%), Naïve Bayes 73% (+/- 15.2%), and KNN 70% (+/- 0%). **C.** Results for data fabricated with the imputation method, mean classification accuracy: Random Forest 90% (+/- 3.4%), GBC 89% (+/- 6.4%), Naïve Bayes 66% (+/- 7.4%), and KNN 56% (+/- 5.3%).

### Machine learning with digit preference

We were unsatisfied with the classification accuracy of the above models. One challenge for machine learning in our data is that the number of features (~17,000) far exceeds the number of samples (75). We therefore explored ways to reduce or transform the feature set, and also to make the feature set more general and broadly applicable. Intrigued by the success of digit frequency methods in the identification of financial fraud [21], we evaluated whether this type of data representation could work for bioinformatics data as well. Therefore, all copy number data was transformed into 20 features, representing the digits 0-9 in the first and second place after the decimal of each gene expression value. While Benford’s Law describes the frequency of the first digit, genomics and proteomics data are frequently normalized or scaled and so the first digit may not be as characteristic. For this reason, our method may be accurately referred to as Benford’s Law inspired or Benford-like. These features were tabulated for each sample to create a new data representation and fed into the exact same machine learning training and testing routine described above. Each of these 20 new features contain decimal values ranging from 0.0 to 1.0 representative of the proportional frequency that digit occurs. For example, one sample’s value in the feature column for the digit 1 may contain the value 0.3. This means that in this sample’s original data the digit 1 occurred in the first position after the decimal place 30% of the time.

In addition to reducing the number of features, converting all features into digit frequencies improves the model’s generality. Machine learning typically cannot handle data where the features are not consistent in number and type. Converting all measured variables to digit frequencies circumvents this problem. For instance, if you had a data set of CNA and transcriptomic data a machine learning model could not train and test on both of these. The features in these datasets would differ in the number of features and what these features represent. But once all information has been converted into digit frequencies the number and type of features are standardized, enabling the model to work any number of different datasets.

In sharp contrast to the models built on the quantitative copy number data, machine learning models which utilized the digit frequencies were highly accurate and showed little variability over the 50 trails (Figure 3). When examining the results of the data fabricated via imputation (both the most sophisticated and most realistic), the models achieved impressively high accuracy. As an average accuracy for the 50 trials, both random forest and the gradient boosting models achieved 100% accuracy. The naïve Bayes model was highly successful with a mean classification accuracy 97%.

**Figure 3 -.**
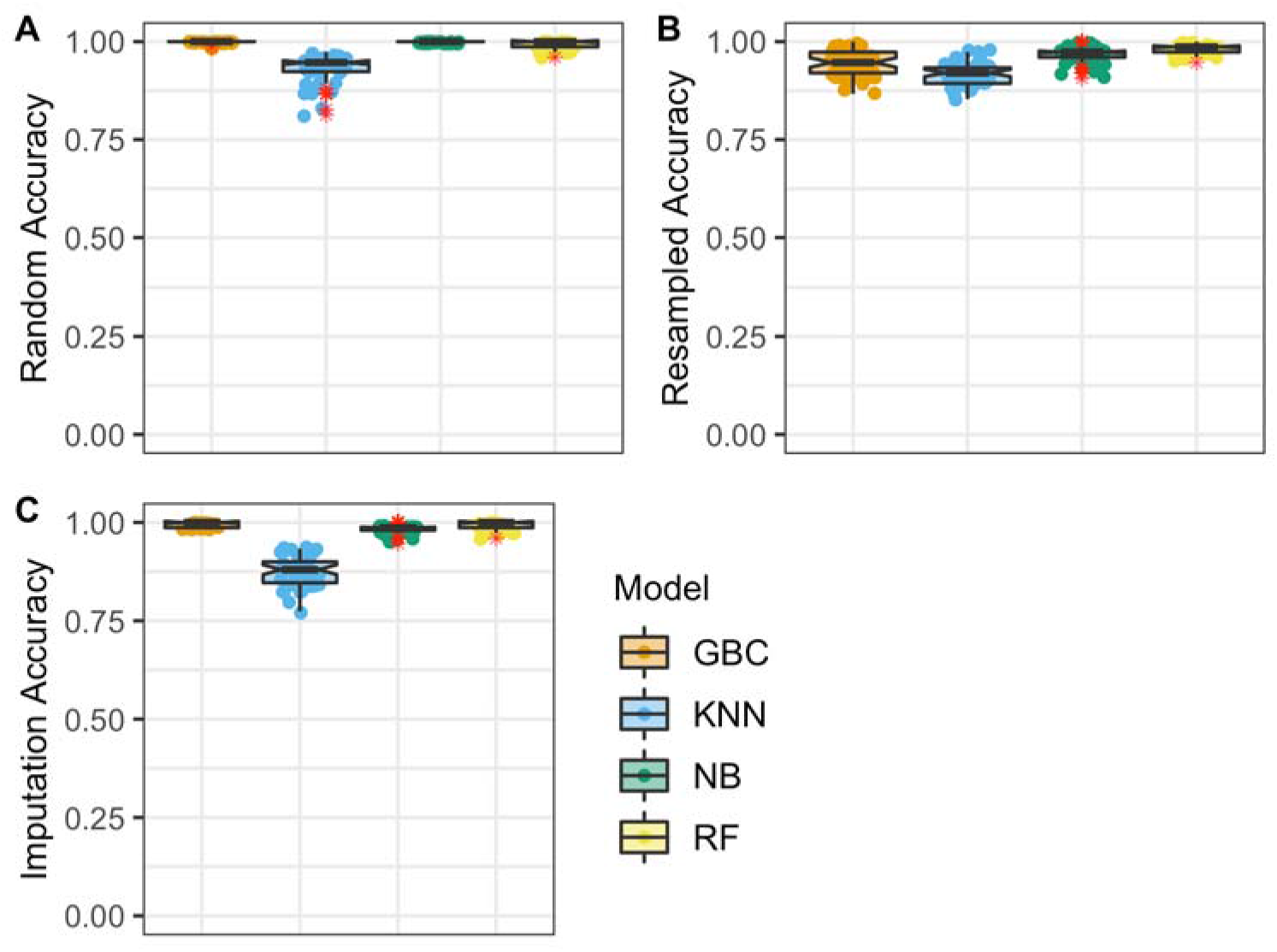
Classifications accuracy using digit frequency data. Fabricated data was mixed with real data and given to four machine learning models for classification. Data shown represents 50 trials for 50 different fabricated dataset mixes. Features in this dataset are the digit frequencies for each sample. **A.** Results for data fabricated with the random method, mean classification accuracy: Random Forest 99% (+/- 1.0%), GBC 100% (+/- 0.2%), Naïve Bayes 100% (+/- 0.0%), and KNN 93% (+/- 3.4%). **B.** Results for data fabricated with the resampling method, mean classification accuracy: Random Forest 98% (+/- 1.3%), GBC 94% (+/- 3.5%), Naïve Bayes 97% (+/- 2.1%), and KNN 92% (+/- 2.8%). **C.** Results for data fabricated with the imputation method, mean classification accuracy: Random Forest 100% (+/- 1.0%), GBC 100% (+/- 0.7%), Naïve Bayes 97% (+/- 1.1%), and KNN 89% (+/- 3.8%).

### Machine learning with limited data

With 17,000 CNA gene measurements, the digit frequencies represent a well sampled distribution. Theoretically, we realize that if one had an extremely limited dataset with CNA measurements for only 10 genes, the sampling of the frequencies for the 10 digits will be poor. To understand how much data is required for a good sampling of the digit-frequencies, we iteratively downsampled our measurements from 17,000 to 10. With the gene-features remaining in each downsample, the digit frequencies were re-calculated. Downsampling was performed uniformly at random without replacement. For each measurement size 100 replicates were run, all with different permutations of the downsamples. Results from this experiment can be seen in Figure 4. The number of gene-features used to calculate digit frequencies does not appear to make a difference at n > 500. In the 100 gene-feature trial, both Naive Bayes and KNN have a significant drop in performance, while the Random Forest and Gradient Boosting model remained relatively unaffected down to approximately 40 features. Surprisingly, these top performing models (GBC and Random Forest) do not drop below 95% accuracy until they have less than 20 gene-features.

**Figure 4 -.**
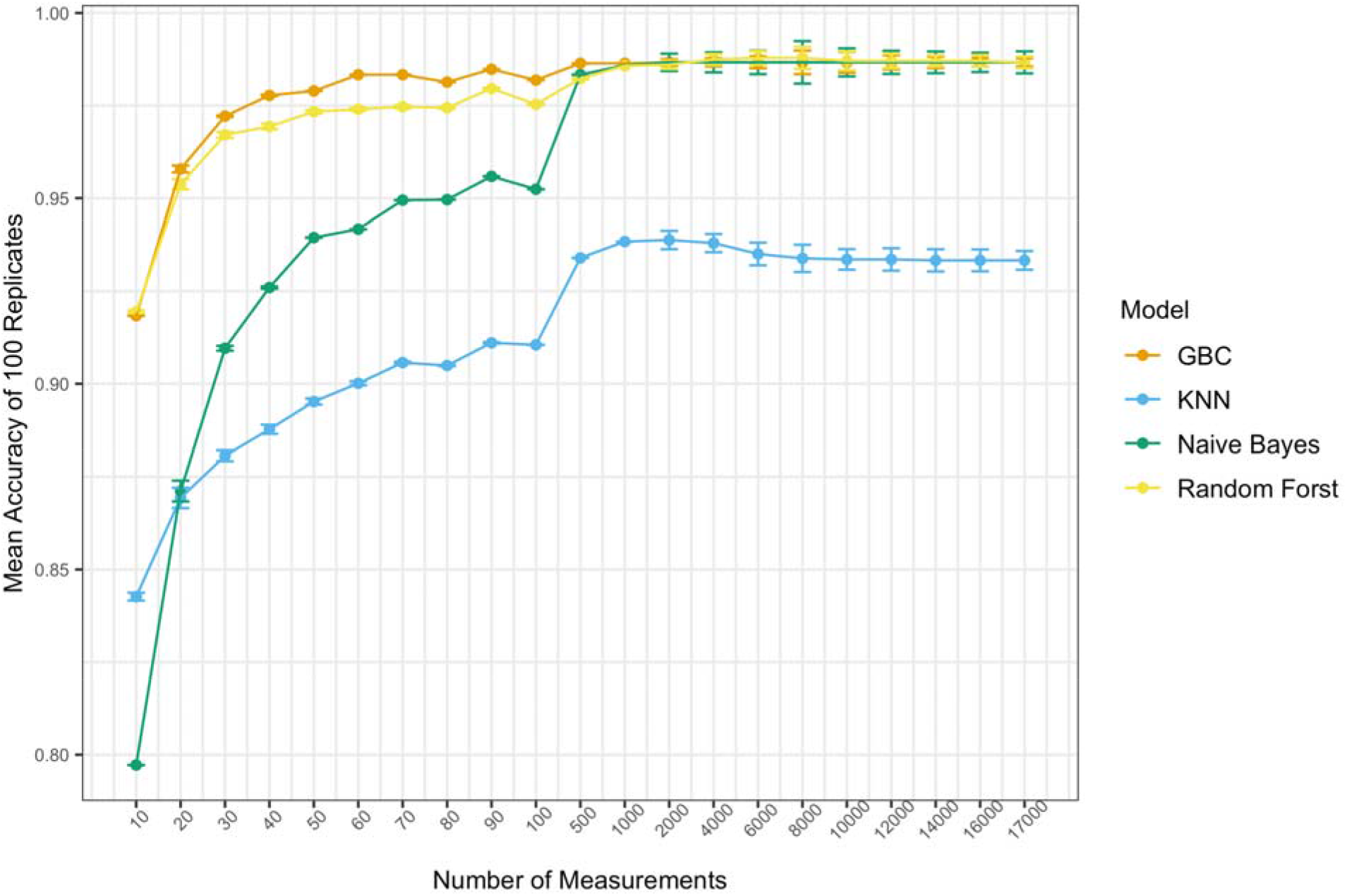
Classifications accuracy vs number of features. The original 17,000 CNA measurements were randomly downsampled incrementally to 10 and converted to digit-frequency training and test features for machine learning models. When 1,000+ measurements are used in the creation of digit-preference features, there appears to be little to no effect on mean accuracy. Below 1,000 Naive Bayes and KNN models begin to lose accuracy quickly. GBC and Random Forest do suffer in accuracy as the number measurements used to generate features lowers but remain above 95% accurate until less than 20 measurements are included.

One hesitation for using machine learning with smaller datasets (i.e. fewer gene-features per data point) is the perceived susceptibility to large variation in performance. As noted, these downsampling experiments were performed 100 times, and error bars representing the standard error are shown in Figure 4. We note that even for the smallest datasets, performance does not noticeably vary between the 100 trials. In fact the standard error for small datasets (e.g. 20 or 30 gene-features) is lower than when there were thousands. Thus we believe that the digit-frequency based models will perform well on both large-scale omics data and also on smaller ‘targeted’ data acquisition paradigms like multiplexed PCR or MRM proteomics.

## Discussion

We present here a proof of concept method for detecting fabrication in biomedical data. Just as has been previously shown in the financial sector, digit frequencies are a powerful data representation when used in combination with machine learning to predict the authenticity of data. Although the data used herein is copy number variation from a cancer cohort, we believe that the Benford-like digit frequency method can be generalized to any tabular numeric data. While multiple methods of fabrication were used, we acknowledge there are more subtle or sophisticated methods. We believe that fraud detection methods, like the models presented herein, could be refined and generalized for broad use in monitoring and oversight.

There is an increasing call for improved oversight and review of scientific data[5,6,16,18], and various regulatory bodies or funding agencies could enforce scientific integrity through the application of these or similar methods. For example, the government bodies charged with evaluating the efficacy of new medicine could employ such techniques to screen large datasets that are submitted as evidence for the approval of new drugs. For fundamental research, publishers could mandate the submission of all data to fraud monitoring. Although journals commonly use software tools to detect plagiarism in the written text, a generalized computational tool focused on data could make data fraud detection equally simple.

## Acknowledgments

This work was supported by the National Cancer Institute (NCI) CPTAC award [U24 CA210972].

## Supplemental Figures

**Supplemental Figure 1 -.**
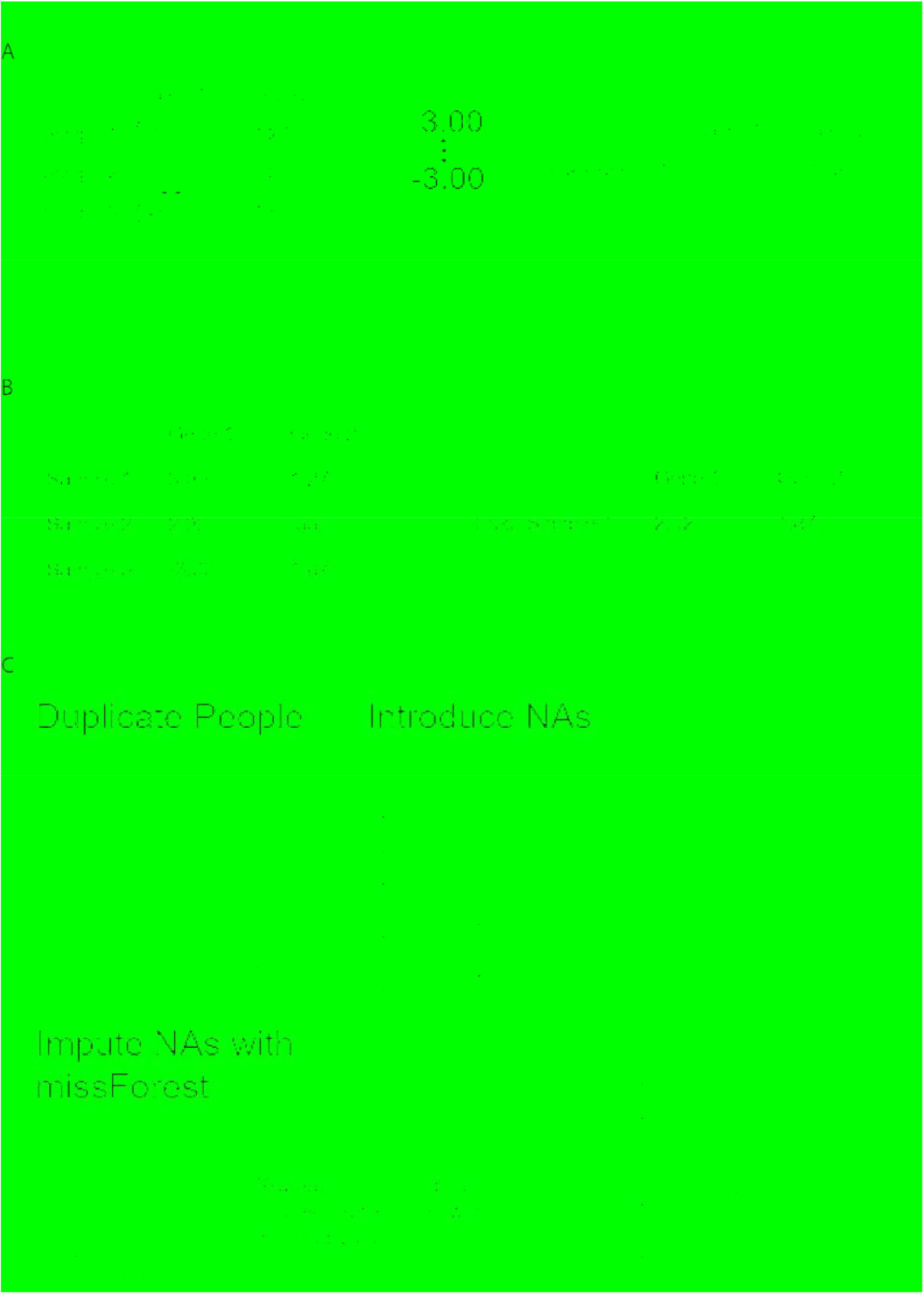
Methods of Data fabrication. (A) The random method of data fabrication identifies the range of observation for a specific locus and then randomly chooses a number in that range. (B) The resampling method chooses values present in the original data. (C) The imputation method iteratively nullifies and then imputes data points from a real sample.

**Supplemental Figure 2 -.**
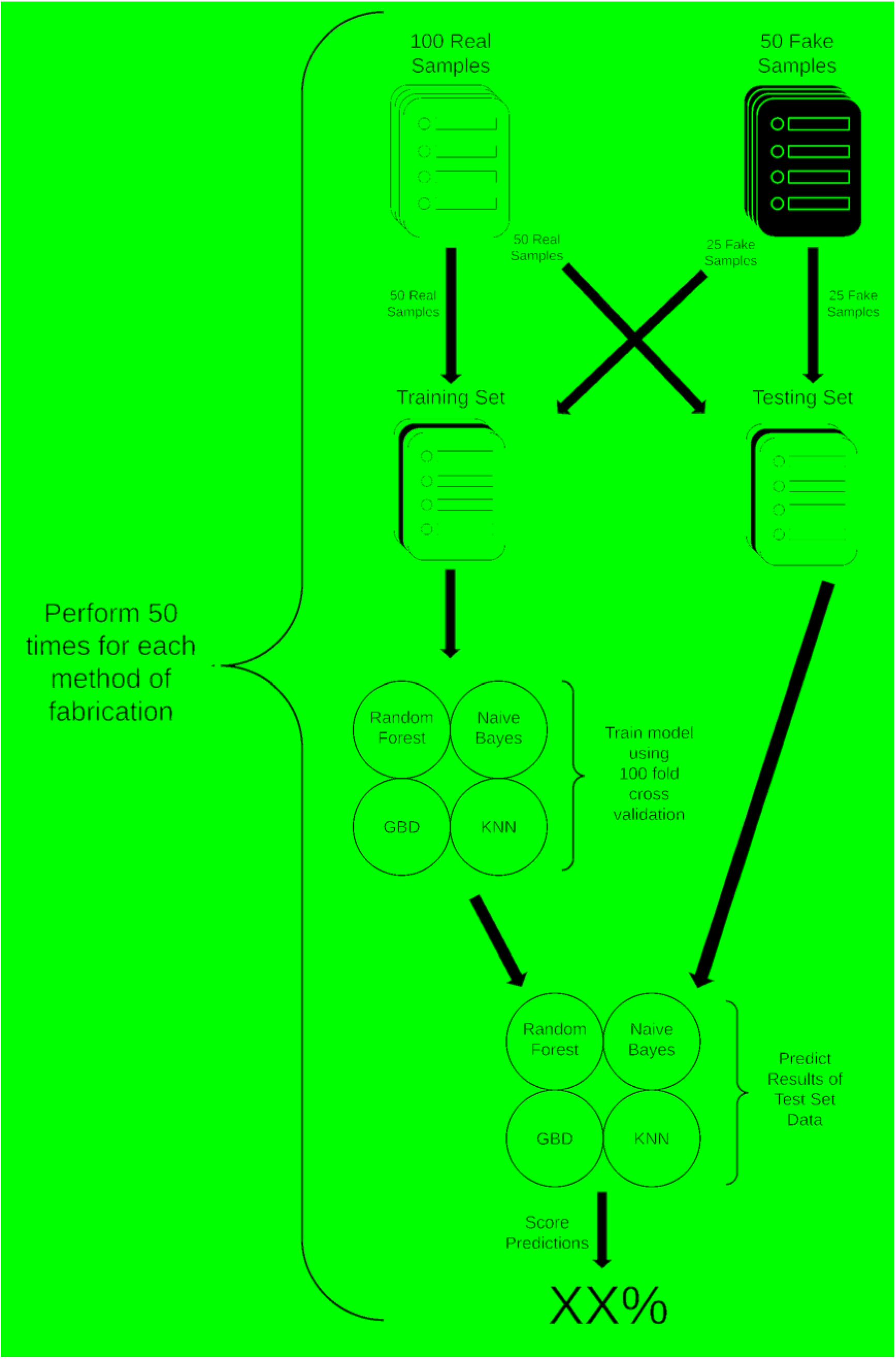
Training and testing overview. After creating 50 fake samples using any one of the three methods of fabrication, the 100 real samples and 50 fake samples were randomly split into a train and test set of equal size and proportions (50 real and 25 fake in each set). The training sets were then used to train various machine learning models using 10-fold cross validation. Next, trained models were used to make predictions on the testing data. Predictions were then scored with total accuracy.

**Supplemental Figure 3 -.**
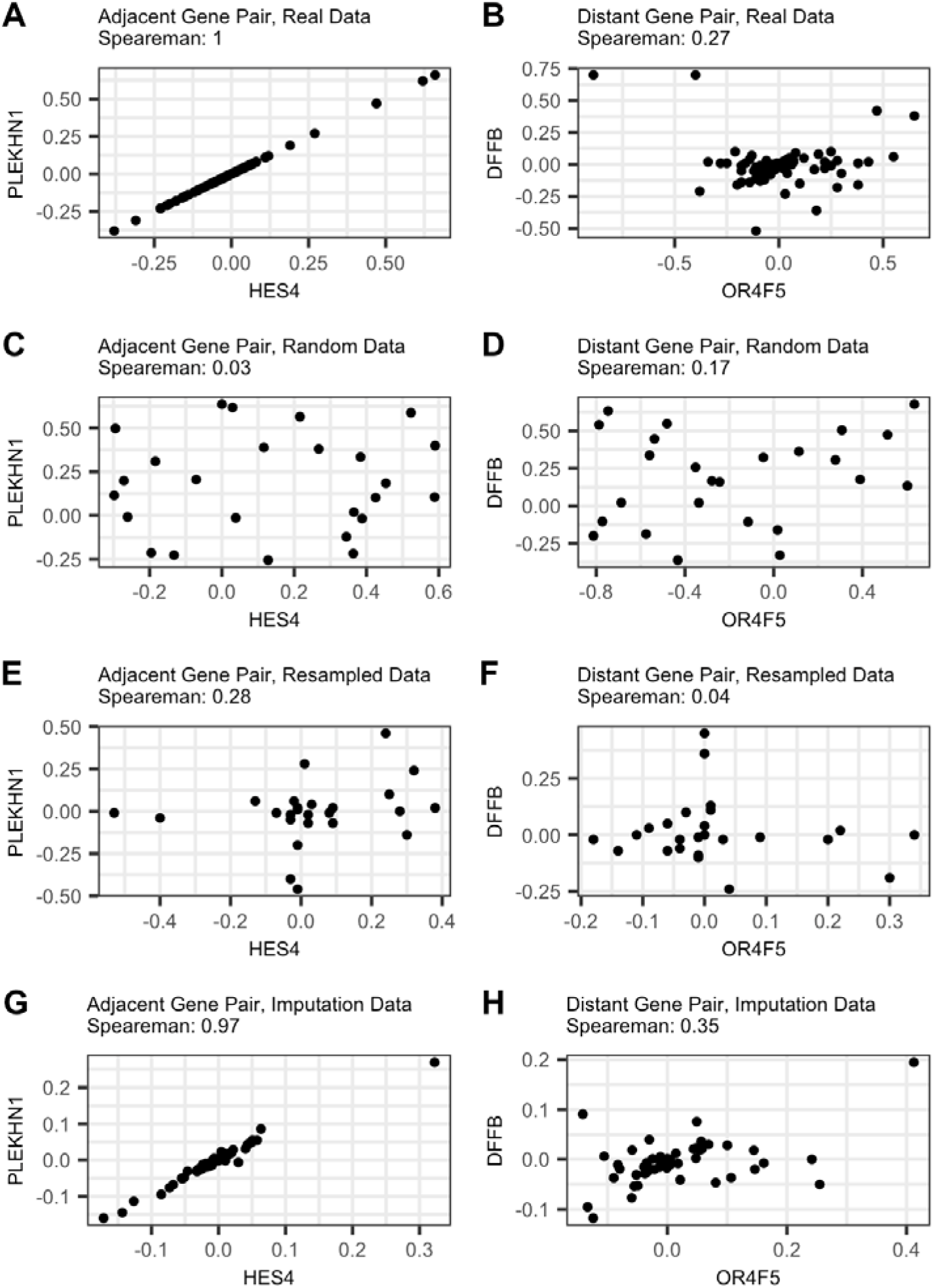
Data relationships in fabricated data. The correlation between pairs of genes is evaluated to determine whether fabrication methods can replicate inter-gene patterns. Plots on the left hand side (A,C,E, and G) display data from two correlated genes *PLEKHN1* and *HES4*, adjacent genes found on 1p36. Plots on the right hand side (B,D,F, and H) display genes *DFFB* and *OR4F5* gene with marginal Spearman correlation in the real data (0.27). The plots reveal that random and resample data have little to no correlation between related genes. Imputation produces data with correlation values that are similar to the original data (0.97 and 0.35, respectively).

## References

1. Burton F. The acquired immunodeficiency syndrome and mosquitoes. Med J Aust. 1989;151: 539–540.

2. Kupferschmidt K. Tide of lies. Science. 2018;361: 636–641.

3. Al-Marzouki S, Evans S, Marshall T, Roberts I. Are these data real? Statistical methods for the detection of data fabrication in clinical trials. BMJ. 2005;331: 267–270.

4. Fanelli D. How many scientists fabricate and falsify research? A systematic review and meta-analysis of survey data. PLoS One. 2009;4: e5738.

5. George SL, Buyse M. Data fraud in clinical trials. Clin Investig. 2015;5: 161–173.

6. Yu L, Miao M, Liu W, Zhang B, Zhang P. Scientific Misconduct and Associated Factors: A Survey of Researchers in Three Chinese Tertiary Hospitals. Account Res. 2020. doi:10.1080/08989621.2020.1809386

7. Blum A, Wang P, Zenklusen JC. SnapShot: TCGA-Analyzed Tumors. Cell. 2018;173: 530.

8. TEDDY Study Group. The Environmental Determinants of Diabetes in the Young (TEDDY) study: study design. Pediatr Diabetes. 2007;8: 286–298.

9. Orwoll E, Blank JB, Barrett-Connor E, Cauley J, Cummings S, Ensrud K, et al. Design and baseline characteristics of the osteoporotic fractures in men (MrOS) study--a large observational study of the determinants of fracture in older men. Contemp Clin Trials. 2005;26: 569–585.

10. Bycroft C, Freeman C, Petkova D, Band G, Elliott LT, Sharp K, et al. The UK Biobank resource with deep phenotyping and genomic data. Nature. 2018;562: 203–209.

11. Barretina J, Caponigro G, Stransky N, Venkatesan K, Margolin AA, Kim S, et al. The Cancer Cell Line Encyclopedia enables predictive modelling of anticancer drug sensitivity. Nature. 2012;483: 603–607.

12. Subramanian A, Narayan R, Corsello SM, Peck DD, Natoli TE, Lu X, et al. A Next Generation Connectivity Map: L1000 Platform and the First 1,000,000 Profiles. Cell. 2017;171: 1437–1452.e17.

13. Caswell J, Gans JD, Generous N, Hudson CM, Merkley E, Johnson C, et al. Defending Our Public Biological Databases as a Global Critical Infrastructure. Front Bioeng Biotechnol. 2019;7: 58.

14. Bik EM, Casadevall A, Fang FC. The Prevalence of Inappropriate Image Duplication in Biomedical Research Publications. MBio. 2016;7. doi:10.1128/mBio.00809-16

15. Knepper D, Fenske C, Nadolny P, Bedding A, Gribkova E, Polzer J, et al. Detecting Data Quality Issues in Clinical Trials: Current Practices and Recommendations. Ther Innov Regul Sci. 2016;50: 15–21.

16. Baigent C, Harrell FE, Buyse M, Emberson JR, Altman DG. Ensuring trial validity by data quality assurance and diversification of monitoring methods. Clin Trials. 2008;5: 49–55.

17. Morrison BW, Cochran CJ, White JG, Harley J, Kleppinger CF, Liu A, et al. Monitoring the quality of conduct of clinical trials: a survey of current practices. Clin Trials. 2011;8: 342–349.

18. Calis KA, Archdeacon P, Bain R, DeMets D, Donohue M, Elzarrad MK, et al. Recommendations for data monitoring committees from the Clinical Trials Transformation Initiative. Clin Trials. 2017;14: 342–348.

19. Benford F, Langmuir I. The Law of Anomalous Numbers. American Philosophical Society; 1938.

20. Cerioli A, Barabesi L, Cerasa A, Menegatti M, Perrotta D. Newcomb-Benford law and the detection of frauds in international trade. Proc Natl Acad Sci U S A. 2019;116: 106–115.

21. Badal-Valero E, Alvarez-Jareño JA, Pavía JM. Combining Benford’s Law and machine learning to detect money laundering. An actual Spanish court case. Forensic Sci Int. 2018;282: 24–34.

22. George SL. Research misconduct and data fraud in clinical trials: prevalence and causal factors. Int J Clin Oncol. 2016;21: 15–21.

23. Stekhoven DJ, Bühlmann P. MissForest--non-parametric missing value imputation for mixed-type data. Bioinformatics. 2012;28: 112–118.

